# Disparate patterns of thermal adaptation between life stages in temperate vs. tropical *Drosophila melanogaster*

**DOI:** 10.1101/120360

**Authors:** Brent L. Lockwood, Tarun Gupta, Rosemary Scavotto

## Abstract

Many terrestrial ectothermic species exhibit limited variation in upper thermal tolerance across latitude. However, these trends may not signify limited adaptive capacity to increase thermal tolerance in the face of climate change. Instead, thermal tolerance may be similar among populations because behavioral thermoregulation by mobile organisms or life stages may buffer natural selection for thermal tolerance. We compared thermal tolerance of adults and embryos among natural populations of *Drosophila melanogaster* from a broad range of thermal habitats around the globe to assess natural variation of thermal tolerance in mobile vs. immobile life stages. We found no variation among populations in adult thermal tolerance, but embryonic thermal tolerance was higher in tropical strains than in temperate strains. Average maximum temperature of the warmest month of the year predicted embryonic thermal tolerance in tropical but not temperate sites. We further report that embryos live closer to their upper thermal limits than adultso—i.e., thermal safety margins are smaller for embryos than adults. F1 hybrid embryos from crosses between temperate and tropical populations had thermal tolerance that matched that of tropical embryos, suggesting dominance of heat-tolerant alleles. Together our findings suggest that thermal selection has led to divergence in embryonic thermal tolerance but that selection for divergent thermal tolerance may be limited in adults. Further, our results suggest that thermal traits should be measured across life stages in order to better predict adaptive limits.

**Impact Summary:** Climate change may threaten the extinction of many ectothermic species, unless populations can evolutionarily adapt to rising temperatures. Natural selection should favor individuals with higher heat tolerances in hotter environments. But recent studies have found that individuals from hot and cold places often have similar heat tolerances. This pattern may indicate that the evolution of heat tolerance is constrained. If this were true, then it would have dire consequences for species persistence under novel thermal conditions.

An alternative explanation for lack of variation in heat tolerance is that mobile organisms don’t need higher heat tolerances to survive in hotter places. The majority of studies have focused on heat tolerance of the adult life stage. Yet, adults in many species are mobile organisms that can avoid extreme heat by seeking shelter in cooler microhabitats (e.g., shaded locations). In contrast, immobile life stages (e.g., insect eggs) cannot behaviorally avoid extreme heat. Thus, mobile and immobile life stages may face different thermal selection pressures that lead to disparate patterns of thermal adaptation across life stages.

Here, we compared heat tolerances of fruit fly adults and eggs (*Drosophila melanogaster*) from populations in temperate North America and tropical locations around the globe. Consistent with previous studies, we found no differences among populations in adult heat tolerance. However, eggs from tropical flies were consistently more heat tolerant than eggs from North American flies. Further, eggs had lower heat tolerance than adults. Consequently, fly eggs in the hotter tropics may experience heat death more frequently than adult flies later in life. This may explain why patterns of divergence in heat tolerance were decoupled across life stages. These patterns indicate that thermal adaptation may be life-stage-specific and suggest that future work should characterize thermal traits across life stages to better understand the evolution of thermal limits.

## Introduction

Extreme temperatures, which may be encountered at the edge of a species’ geographic range (Hilbish *et al.* 2010) or episodically during the hottest or coldest days of the year (Hoffmann 2010; Kingsolver, Diamond & Buckley 2013; Dowd, King & Denny 2015; Buckley & Huey 2016), can cause populations to experience mortality (Helmuth *et al.* 2002; Denny, Miller & Harley 2006) and ultimately lead to thermal adaptation (Lenski & Bennett 1993; Mongold, Bennett & Lenski 1999; Hangartner & Hoffmann 2015). However, recent work suggests that thermal adaptation of upper thermal limits might be evolutionarily constrained (Hoffmann, Chown & Clusella-trullas 2013; Schou *et al.* 2014; Hangartner & Hoffmann 2015; Kristensen *et al.* 2015; van Heerwaarden, Kellermann & Sgrò 2016), such that the evolution of increased heat tolerance might be a relatively slow process that cannot occur over short evolutionary timescales (Kellermann *et al.* 2012). If this is the case, global climate change, which has led to rapid increases in mean temperatures and the frequency of extreme thermal events (Katz & Brown 1992; Meehl *et al.* 2000; Cai *et al.* 2014), may cause shifts in geographic distributions (Rank & Dahlhoff 2002; Burrows *et al.* 2011; Thomas *et al.* 2012; Sunday, Bates & Dulvy 2012) as populations may not be able to adapt fast enough to persist in hotter environments (Jezkova & Wiens 2016).

But thermal adaptation depends on the strength of selection (Bennett, Lenski & Mittler 1992; Rudolph *et al.* 2010), and studies that focus on thermal tolerance of mobile organisms or life stages may overestimate the degree to which these organisms encounter thermal selection in nature. In other words, thermal safety margins—i.e., the difference between upper thermal limits and maximum habitat temperature—may be larger than predicted because thermal environmental heterogeneity allows mobile organisms to avoid thermal extremes via behavioral thermoregulation (Dillon *et al.* 2009; Gunderson & Leal 2012; Buckley, Ehrenberger & Angilletta 2015; Llewelyn *et al.* 2016; Munoz *et al.* 2016). To date, there have been relatively few studies that examine thermal tolerance in immobile organisms or life stages, particularly in the terrestrial realm (Angilletta *et al.* 2013; MacLean *et al.* 2016), and immobile organisms may represent ideal study systems to investigate the evolutionary potential of thermal tolerance. In support of this conjecture, broad scale patterns of thermal tolerance are more tightly correlated with habitat temperatures in marine systems than in terrestrial systems (Sunday, Bates & Dulvy 2011), perhaps due to the more limited range of thermal microhabitats in the marine realm (Denny *et al.* 2011) that makes behavioral thermoregulation a less effective buffering mechanism.

Here we sought to compare adult and embryonic heat tolerance among populations of fruit flies, *Drosophila melanogaster*, from a broad range of thermal habitats across the world to ascertain the degree to which thermal selection has shaped the evolution of thermal tolerance across immobile vs. mobile life stages. Adult thermal tolerance has been extensively studied in natural populations of *D. melanogaster* (Bettencourt *et al.* 2002; Hoffmann & Weeks 2007; Adrion, Hahn & Cooper 2015; Buckley & Huey 2016), but to a large extent the thermal physiology of the early embryonic life stage of *D. melanogaster* has not been characterized in natural populations (Sgro *et al.* 2010; Overgaard, Kearney & Hoffmann 2014; Kristensen *et al.* 2015). Studies of laboratory-bred *D. melanogaster* have shown that early embryos (0 – 2 hours post-fertilization) are more thermally sensitive than later stages (Walter, Biessmann & Petersen 1990), perhaps due to the reduced heat-shock response in early embryos (Graziosi *et al.* 1980; Welte *et al.* 1993). Thus, we compared heat tolerance of adults and early stage embryos to determine whether or not differences in thermal sensitivity, as well as mobility, lead to different patterns of thermal adaptation across life stages. The thermal environment of *D. melanogaster* can change rapidly (+18°C h^-1^) and reach extreme values (> 40°C) (Feder, Blair & Figueras 1997; Terblanche *et al.* 2011). Therefore, we designed our thermal stress experiments to mimic sudden (acute) changes in temperature that are characteristic of the variable thermal environments that flies experience in nature (Terblanche *et al.* 2011). We report higher embryonic thermal tolerance in tropical (hotter) vs. temperate (cooler) populations but no difference in adult thermal tolerance, and thus we demonstrate that selection for thermal tolerance likely varies across life stages. Moreover, our data suggest that there is significant adaptive variation for upper thermal tolerance in natural populations in the earliest and most thermally sensitive life stage.

## Materials and methods

### Fly strains

We obtained 20 isofemale genetic lines that were collected from temperate locations in the USA as a generous gift from B.S. Cooper and K.L. Montooth: 6 lines from Raleigh, NC (NC); 6 lines from Beasley Orchard, IN (IN); and 8 lines from East Calais, VT (VT). These lines were established by single female founders whose progeny were subsequently inbred for several generations to isogenize the genetic variability within each line, and thereby minimize the potential for lab evolution (Cooper, Hammad & Montooth 2014). These temperate North American lines have been maintained at controlled densities of 50 to 100 adults per vial since their establishment. We obtained 5 isofemale lines from the Drosophila Species Stock Center at the University of California, San Diego that were collected from tropical locations around the world: 1 line each from Accra, Ghana (GH); Mumbai, India (MU); Guam, USA (GU); Chiapas, Mexico (CH); and Monkey Hill, St. Kitts (SK). Stocks from the UCSD Stock Center were also established by single female founders, as described above for the North American isofemale lines, and have been maintained at controlled densities since their establishment. Geographic coordinates of collection locations are shown in Table 1 and stock numbers and collection dates of isofemale lines are provided in Supplementary Table S1. We maintained flies under common-garden conditions on cornmeal-yeast-molasses medium at 25°C on a 12:12 light cycle for at least two generations prior to measuring thermal tolerance.

**Table 1:** Collection site locations, regions, climate zones, and maximum habitat temperatures of the warmest month of the year (T_max_) from 1950-2000 (WorldClim; Hijmans *et al.* 2005).

| Collection Locale | Lat. (°N) | Long. (°E) | Region | Climate Zone | $T_{\max}$ (°C) |
| --- | --- | --- | --- | --- | --- |
| East Calais, Vermont, USA (VT) | 44.4 | -72.4 | North America | North Temperate | 25.7 |
| Beasley Orchard, Indiana, USA (IN) | 39.8 | -86.5 | North America | North Temperate | 29.6 |
| Raleigh, North Carolina, USA (NC) | 35.8 | -78.7 | North America | North Temperate | 31.7 |
| Chiapa de Corzo, Chiapas, Mexico (CH) | 16.7 | -93.0 | Central America | Tropics | 34.1 |
| Monkey Hill, St. Kitts (SK) | 17.3 | -62.7 | Caribbean | Tropics | 30.3 |
| Accra, Ghana (GH) | 5.6 | -0.2 | West Africa | Tropics | 32.0 |
| Mumbai, India (MU) | 19.1 | 72.9 | Western India | Tropics | 33.2 |
| Guam, USA (GU) | 13.4 | 144.8 | Oceania | Tropics | 30.6 |

### Adult thermal tolerance (LT_50_) and critical thermal maximum (CT_max_)

We assayed thermal tolerance (LT_50_) of adult flies by scoring the number of flies surviving after exposure to a 45-minute heat treatment across a range of temperatures, from 36°C to 42°C. 30 minutes prior to heat treatment, 40 adult flies (3 to 5 day-old males and females of equal numbers) were transferred to empty glass vials (25 x 95 mm with Flugs closures, Genesee Scientific, San Diego, CA) and returned to an incubator at 25°C. Vials were then partially submerged in a water bath (1 cm below the top of the vial) and heat shocked for 45 minutes. We monitored the heat ramping rate in these heat treatments with a thermocouple (Omega Engineering, Inc., Norwalk, CT) suspended inside an adjacent empty vial. These heat treatments produced linear heat ramps that were consistent across all temperatures, with an average (± standard deviation) rate of change of +0.6 ± 0.01°C min^-1^. This rate of increase is within the range of measured rates of change in nature (Feder *et al.* 1997; Terblanche *et al.* 2011). Flies were then gently transferred to a food vial, and survival was scored after 24 h of recovery at 25°C. We replicated our treatments across 3 replicate vials at each of four temperatures (36°C, 38°C, 40°C, and 42°C) for each isofemale line (n = 40 flies x 3 vials x 4 temperatures = 480 adults per isofemale line). We scored LT_50_ as the temperature at which 50% of the adults did not recover from heat stress via a least-squares regression model of the logistic equation. We conducted these curve fitting analyses in GraphPad Prism 7 for Mac OS X (GraphPad Software, La Jolla, CA).

To more fully describe adult thermal tolerance among our isofemale lines, we also measured the temperature at which flies incurred a loss of motor response along a heat rampo—i.e., the critical thermal maximum (CT_max_). While previous studies have reported similar values of LT_50_ and CT_max_ in *D. melanogaster* (Huey, Partridge & Fowler 1991; Gilchrist, Huey & Partridge 1997), different thermal tolerance assay methods have been shown to affect the extent to which populations of *D. melanogaster* populations exhibit clinal variation in thermal tolerance (Sgro *et al.* 2010). Thus, we sought to compare both adult LT_50_ and CT_max_ among populations in order to account for potential bias that may be inherent to the assay method. 3 to 5 day-old adult male flies were individually placed into glass vials with rubber stoppers, submerged in a water bath at 25°C, and exposed to a heat ramp of +0.1°C min^-1^. We chose this rate of temperature increase based on previously published studies that measured CT_max_ in *Drosophila* (Chown *et al.* 2009; Sgro *et al.* 2010; Kellermann *et al.* 2012) and to mimic the variable thermal environments that flies encounter in nature (Terblanche *et al.* 2011). Flies were regularly checked for responsiveness along the heat ramp by gently tapping the vial, and the temperature at which a fly lost the ability to move was recorded. We scored CT_max_ for each genotype via a least-squares regression model of the logistic equation among 10 flies per genotype and extrapolated CT_max_ from the inflection points of the logistic curves. We conducted these curve fitting analyses in GraphPad Prism 7.

### Embryonic thermal tolerance (LT_50_)

We assayed embryonic thermal tolerance (LT_50_) by measuring survival (hatching success) of early stage embryos, 0 to 1 h post-fertilization, exposed to a 45-minute heat treatment across a range of temperatures, from 25°C to 42°C. We did not assay CT_max_ for embryos because embryos do not possess behavioral characteristics that would permit the assessment of thermal tolerance via loss of motor activity. We designed our heat treatments to mimic sudden increases in temperature that frequently occur in nature where the temperature of necrotic fruit can increase rapidly on hot days (Feder *et al.* 1997; Terblanche *et al.* 2011). 3 to 5 day-old adult flies were allowed to mate and lay eggs on grape juice agar plates (60 x 15 mm) for 1 h at 25°C. Egg plates were then wrapped in Parafilm, submerged in a water bath, and heat shocked for 45 minutes. We monitored the heat ramping rate in these treatments via a thermocouple (Omega Engineering, Inc.) placed at the surface of the egg plate media. These heat treatments produced heat ramps that were similar to those of the adult LT_50_ assays, with an average (± standard deviation) rate of temperature change of +0.57 ± 0.3°C min^-1^. The higher variance in ramping rates among the egg heat treatments, compared to the relatively low variance among the adult assays, was likely due to the presence of the agar in the egg plates, which varied in thickness between 5 and 10 mm. These rates of increase are within the range of measured rates of change of necrotic fruit in nature (Feder *et al.* 1997).

Following heat shock, 20 eggs were transferred on a piece of grape juice agar to fresh food vials and placed at 25°C. Hatching success was scored as the proportion of larvae that successfully hatched by 48 h. We conducted 4 to 6 replicate treatments at each of 9 temperatures (25°C, 28°C, 30°C, 32°C, 34°C, 36°C, 38°C, 40°C, and 42°C) for each isofemale line (n = 20 embryos x 4 replicates x 9 temperatures = 720 embryos per isofemale line). We used these data to calculate the lethal temperature at which 50% of the embryos failed to hatch (LT_50_) via a least-squares regression model of the logistic equation. In our logistic model, we allowed the y-intercept to vary between 0 and 1 and extrapolated the LT_50_ from the inflection point of the logistic curve fit. This approach allowed us to infer thermal tolerance independently from other confounding factors that may influence the measurement of hatching success, such as the presence of unfertilized eggs. We conducted these curve fitting analyses in GraphPad Prism 7.

### Statistical comparisons of thermal tolerance, thermal safety margins, and maternal effects

We compared adult (LT_50_) and embryonic (LT_50_) thermal tolerances among temperate sites (VT, IN, and NC) and all tropical sites pooled together (CH, SK, GH, MU, and GU) with ANOVA. This ANOVA design allowed us to (1) assess variation within and among North American populations to test for clinal variation in North America and (2) compare variation within and between North America vs. the tropics to test for consistent differences between temperate and tropical regions. Pairwise differences were assessed with Tukey’s multiple comparison post-hoc test.

We calculated thermal safety margins as the difference between thermal tolerance (adult LT_50_ or embryo LT_50_) and maximum temperature of the warmest month (T_max_) at each site. We downloaded T_max_ estimates from the WorldClim database (Hijmans *et al.* 2005) (www.worldclim.org) that corresponded to the GPS coordinates of the collection sites of each population (see Table 1). These T_max_ estimates are based on climate data from the years 1950 to 2000. Fine-scale spatial temperature data are not available for these collection sites, but while T_max_ may not perfectly match the thermal environment experienced by flies, variation in T_max_ should reflect relative differences in the thermal environments among locations. In addition, previous studies have shown T_max_ to be a significant predictor of upper thermal limits in *Drosophila* (Kellermann *et al.* 2012). We assessed the main effects of region (temperate vs. tropical), life stage (adult vs. embryo), and their interaction on thermal safety margins via a 2-way ANOVA. Least-squares linear regression was used to assess the relationship between thermal tolerance and T_max_. ANCOVA was used to assess the difference in slopes of regression lines fit to data from temperate vs. tropical sites.

We tested for the potential role of maternal effects in conferring heat tolerance to tropical embryos by conducting reciprocal crosses between the two parental strains that had the highest and lowest LT_50_, Chiapas, MX (CH) and Vermont, USA strain #12 (VT-12), respectively, and measured thermal tolerance of F1 progeny. At this stage of development (0-1 h-old), early embryos have inactive gene transcription and thus their physiology is predicted to depend on maternal factors, such as mRNAs and proteins, loaded into eggs (Tadros & Lipshitz 2009; Blythe & Wieschaus 2015). We used logistic models to fit the hatching success data, as described above, and compared LT_50_s of the parental strains and their F1 progeny by an extra sum-of-squares F-test of the extrapolated LT_50_s. We conducted these analyses in GraphPad Prism 7.

## Results

### Thermal tolerance and thermal safety margins across life stages

We found no difference in adult thermal tolerance among all sites (Figs. 1A and 1B; ANOVA, *F*_*3,20*_ = 0.3134, *P* = 0.8155), with an overall mean LT_50_ (± 95% C.I.) of 39.84 ± 0.12°C. We also did not observe any difference among collection sites in adult thermal tolerance as measured by CT_max_ (Fig. S1; ANOVA, *F*_*3,9*_ = 2.378, *P* = 0.1375). Adult CT_max_ values were slightly lower than LT_50_ values, with an overall mean (± 95% C.I.) of 38.77 ± 0.52°C (Fig. S2). This lower value of CT_max_ may have been due to multiple factors, including the slower ramping rate of the CT_max_ experiments, the thermal sensitivity of locomotor activity, or the fact that we assayed CT_max_ only for males whereas females were included in our assay of LT_50_.

**Figure 1.**
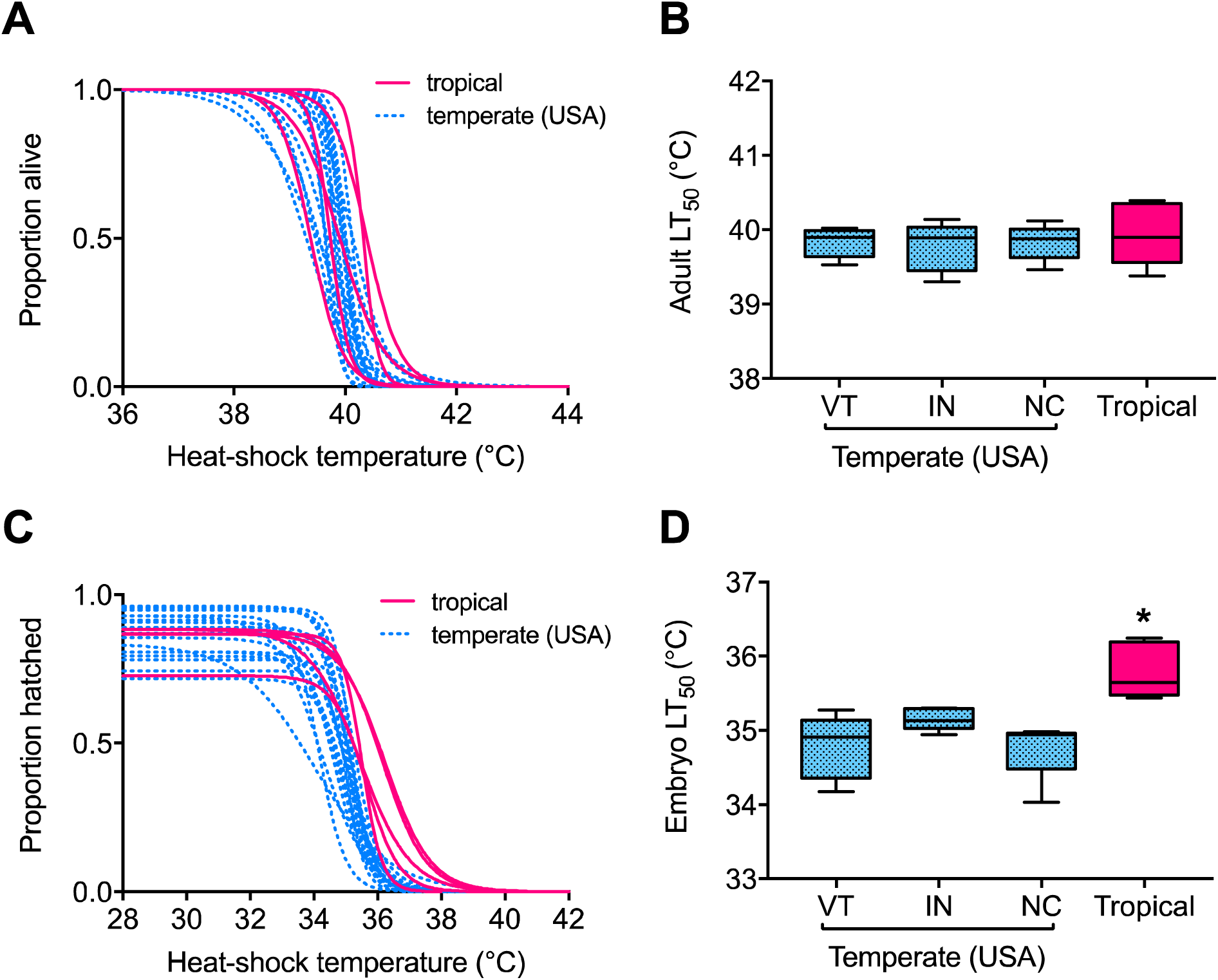
Flies from different populations around the world exhibited differences in embryonic thermal tolerance but not adult thermal tolerance. (A) Proportion of adult flies that survived after heat shock (45 min at indicated temperature; see Methods for rate of temperature change). Tropical lines are indicated in solid pink. Temperate lines are indicated in dotted blue. (**B**) Adult LT_50_ was consistent across all populations (ANOVA, *F*_*3,20*_ = 0.3134, *P* = 0.8155). LT_50_ was extrapolated from the survival curves in A. Boxes indicate upper and lower quartiles, whiskers extend to maximum and minimum values, and horizontal lines indicate the medians. (**C**) Proportion of eggs that successfully hatched following heat shock (45 min at indicated temperature). Tropical lines are indicated in solid pink. Temperate lines are indicated in dotted blue. (**D**) Embryonic thermal tolerance (LT_50_) was higher in tropical lines than temperate lines (ANOVA, *F*_*3,20*_ = 10.16, *P* = 0.0003; Tukey s test, VT vs. IN, *q* = 2.428, *P* = 0.3416, VT vs. NC, *q* = 0.4268, *P* = 0.9902, IN vs. NC, *q* = 2.666, *P* = 0.2656, tropical vs. VT, *q* = 6.909, *P* = 0.0005, tropical vs. IN, *q* = 4.04, *P* = 0.0444, tropical vs. NC, *q* = 4.04, *P* = 0.0005). LT_50_ was extrapolated from the survival curves in C. Boxes and whiskers drawn as in B. \**P* < 0.05.

Embryonic thermal tolerance (LT_50_) did not differ among the three temperate sites but was significantly higher in tropical vs. temperate embryos (Figs. 1C and 1D; ANOVA, *F*_*3,20*_ = 10.16, *P* = 0.0003; Tukey’s test, VT vs. IN, *q* = 2.428, *P* = 0.3416, VT vs. NC, *q* = 0.4268, *P* = 0.9902, IN vs. NC, *q* = 2.666, *P* = 0.2656, tropical vs. VT, *q* = 6.909, *P* = 0.0005, tropical vs. IN, *q* = 4.04, *P* = 0.0444, tropical vs. NC, *q* = 4.04, *P* = 0.0005). Overall, tropical embryos were more heat tolerant; the average LT_50_ was approximately 1°C higher in tropical embryos (35.8 ± 0.45°C) than in temperate embryos (34.88 ± 0.18°C). There was no significant relationship between adult LT_50_ and embryo LT_50_ for either temperate (Fig. S2; Least-squares linear regression, *R*^*2*^ = 0.015, *y* = -0.1973*x* + 42.73) or tropical lines (Fig. S2; Least-squares linear regression, *R*^*2*^ = 0.09, *y* = 0.2664*x* + 25.15).

Thermal safety marginso—i.e., the difference between thermal tolerance (CT_max_ or LT_50_) and maximum habitat temperature (T_max_)—were consistently smaller for embryos than adults. This pattern was consistent across regions (temperate and tropical) (Fig. 2; ANOVA, main effect of life stage, *F*_*1,45*_ = 26.19, *P* < 0.0001), however thermal safety margins were smaller in both life stages for tropical than for temperate sites (Fig. 2; ANOVA, main effect of region, *F*_*1,45*_ = 10.58, *P* = 0.0027, life stage x region interaction, *F*_*1,45*_ = 0.1745, *P* = 0.6782).

**Figure 2.**
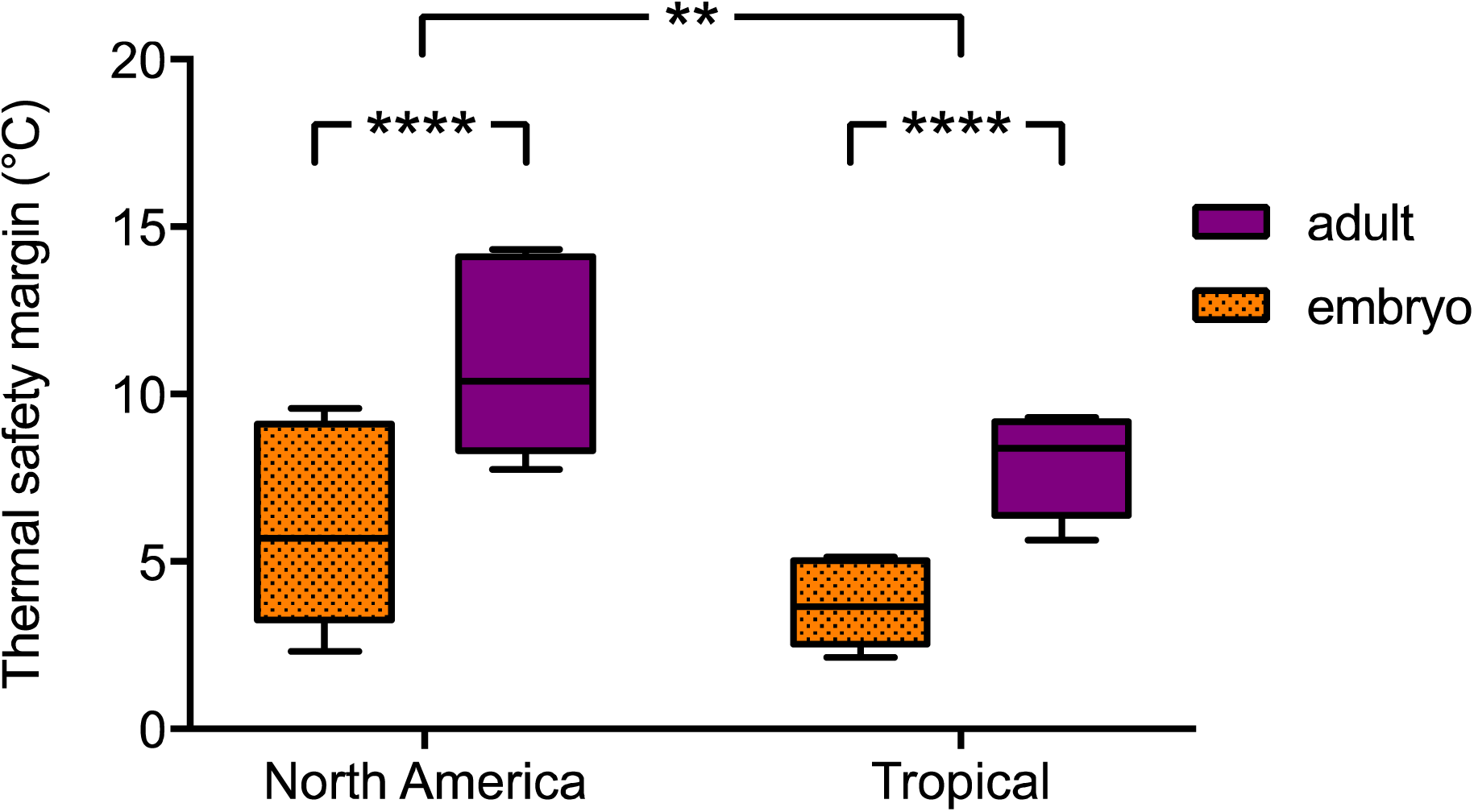
Thermal safety margin differs by life stage and geographic region. (**A**) Thermal safety margins were smaller for embryos than adults and smaller in the tropics than temperate sites (ANOVA, main effect of life stage, *F*_*1,45*_ = 26.19, *P* < 0.0001, main effect of region, *F*_*1,45*_= 10.58, *P* = 0.0027, life stage x region interaction, *F*_*1,45*_= 0.1745, *P* = 0.6782).Boxes indicate upper and lower quartiles, whiskers extend to maximum and minimum values, and horizontal lines indicate the medians. \*\**P* < 0.01, \*\*\*\**P* < 0.0001.

### Maximum habitat temperature and thermal tolerance

Maximum temperature of the warmest month (i.e., maximum habitat temperature or T_max_) spanned a range of 8.4°C among all sites, from 25.7°C in Vermont, USA (VT) to 34.1°C in Chiapas, MX (CH) (Table 1). Previous studies have shown T_max_ to be positively correlated with adult heat tolerance (CT_max_) among many species of *Drosophila* (Kellermann *et al.* 2012); however, our populations of *D. melanogaster* showed no significant relationship between adult heat tolerance (LT_50_) and T_max_ in either temperate (Fig. S3; Least-squares linear regression, *R*^*2*^ = 0.004, *y* = -0.0005*x* + 39.83) or tropical regions (Fig. S3; Least-squares linear regression, *R*^*2*^ = 0.14, *y* = 0.098*x* + 36.82). The embryonic life stage exhibited a different pattern from the adults, and the relationship between embryonic heat tolerance and T_max_ was distinct between temperate and tropical regions (Fig. 3; ANCOVA, *F*_1,4_ = 10.26, *P* = 0.0328). Among temperate populations there was a 6°C range in T_max_, but this produced no correlated response in the thermal tolerance of embryos (Fig. 3; Least-squares regression, *R*^*2*^ = 0.0015, *P* = 0.9751, *y* = 0.00282*x* + 34.82). But among tropical populations, the approximate 4°C range in T_max_ corresponded to a positive relationship between embryonic thermal tolerance and T_max_ (Fig. 3; Least-squares regression, *R*^*2*^ = 0.9478, *P* = 0.0051, *y* = 0.2199*x* + 28.75).

**Figure 3.**
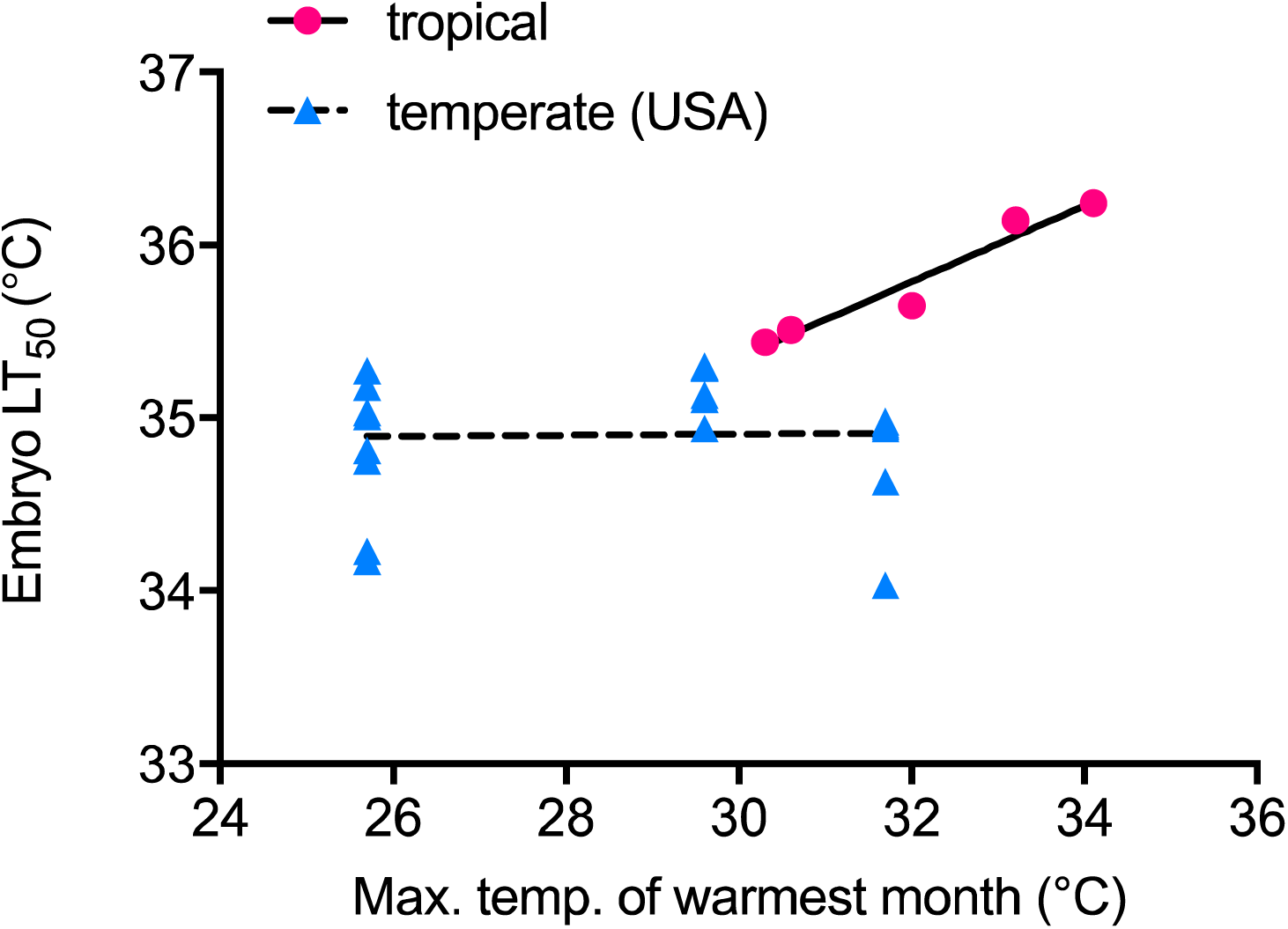
Embryonic thermal tolerance and maximum habitat temperature (T_max_) by region. Embryonic thermal tolerance was positively correlated with T_max_ among tropical populations (Least-squares regression, *R*^*2*^ = 0.9478, *P* = 0.0051, *y* = 0.2199*x* + 28.75) but not temperate populations (Least-squares regression, *R*^*2*^ = 0.0015, *P* = 0.9751, *y* = 0.00282*x* + 34.82). Tropical genotypes are indicated in pink circles, with a solid black regression line fit. Temperate genotypes are indicated in blue triangles, with a dashed black regression line fit.

### Embryonic thermal tolerance in F1 progeny from Chiapas x Vermont

Offspring from reciprocal genetic crosses between the most heat tolerant tropical genotype (CH) and the least heat tolerant temperate genotype (VT-12) had thermal tolerances that closely resembled that of the heat tolerant CH genotype, regardless of the direction of the cross (Fig. 4), suggesting dominance of heat tolerant alleles and no significant maternal effect. Embryonic LT_50_s of F1 progeny of both crosses (CH♀ x VT♂ = 35.83°C and VT♀ x CH♂ = 35.80°C) were statistically indistinguishable from the LT_50_ of CH (36.24°C) but significantly higher than the LT_50_ of VT-12 (34.23°C; Fig. 4; Logistic model, Extra sum-of-squares F-test on lower LT_50_ of VT-12, *F3,166* = 6.695).

**Figure 4.**
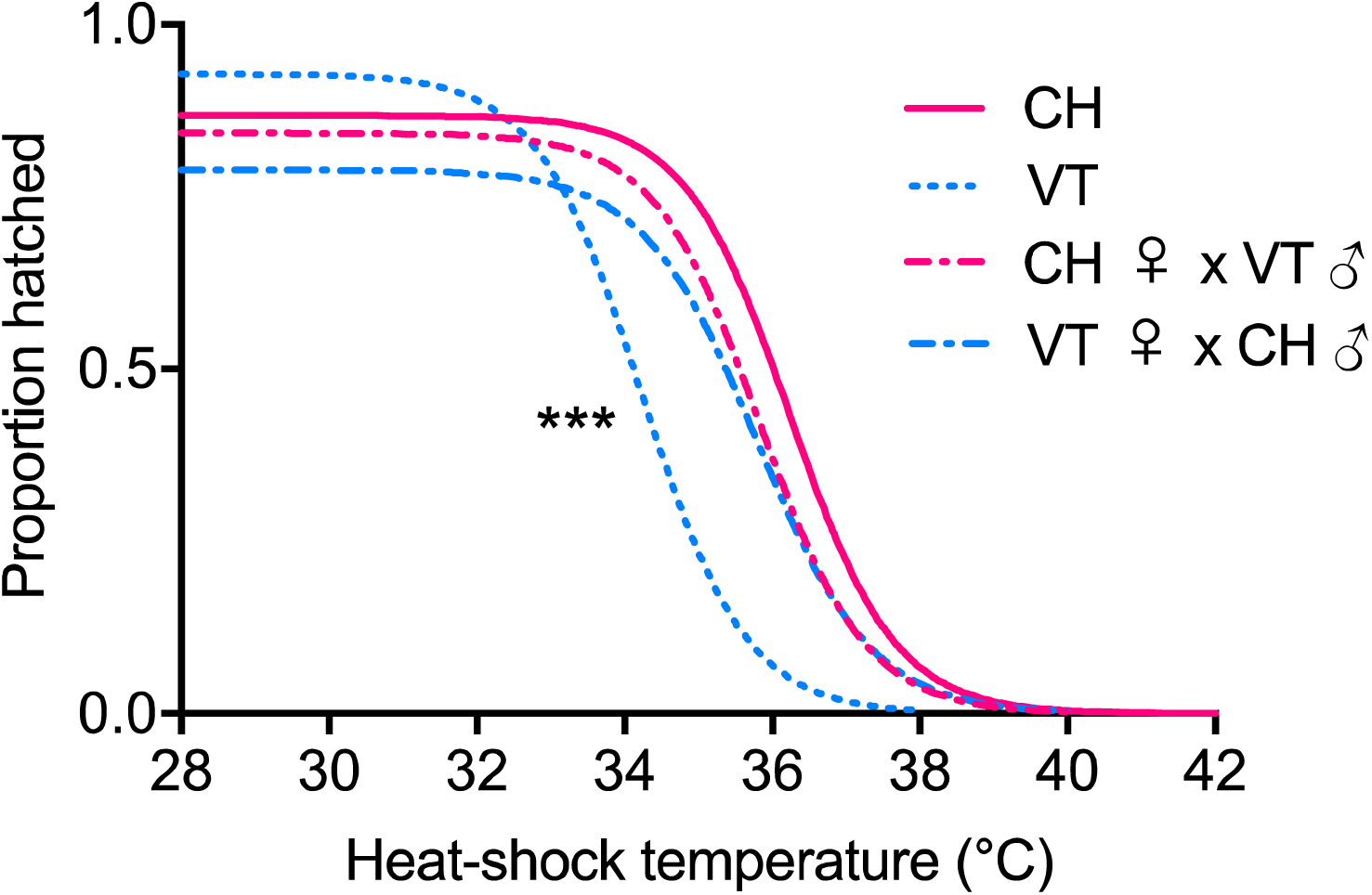
F1 progeny from tropical x temperate parents have high embryonic heat tolerance. Proportion of eggs that successfully hatched following heat shock (45 min at indicated temperature) among two parental genotypes that had the highest and lowest embryonic LT_50_ of all strains in this study, CH (Chiapas, Mexico) and VT-12 (Vermont, USA), respectively, along with F1 progeny from reciprocal crosses of these two parental lines, CH♀ x VT♂ and VT♀ x CH♂ (♀ = dam; ♂ = sire). Note that VT-12 is labeled VT in the legend. LT_50_: CH = 36.24°C, VT-12 = 34.23°C, CH ♀ x VT♂ = 35.83°C, VT♀ x CH♂ = 35.80°C (Logistic model, Extra sum-of-squares F-test on lower LT_50_ of VT-12, *F3,166* = 6.695, \*\*\**P* = 0.0003).

## Discussion

Despite the potential for thermal adaptation across the broad range of thermal habitats represented in this study, our data suggest that natural selection on thermal tolerance does not act equally across life stages in *D. melanogaster*. Rather, we provide evidence of adaptive variation in upper thermal limits in the thermally sensitive and immobile embryonic life stage but not in the more thermally tolerant and mobile adult stage. This is perhaps not surprising, given that lower thermal tolerance in early embryos translates into smaller thermal safety margins. Thus, we predict that embryos encounter lethal temperatures more frequently than adults, particularly because embryos lack the ability to behaviorally avoid thermally stressful conditions, and this likely drives divergence in embryonic thermal tolerance between temperate North American and tropical populations.

Recent estimates of divergence in adult thermal tolerance among populations of *D. melanogaster* have brought into question the degree of adaptive potential in upper thermal limits in this species, as comparisons of populations across latitude have yielded mixed results depending on assay methods (Sgro *et al.* 2010) and the laboratory in which thermal tolerance was measured (Hoffmann, Anderson & Hallas 2002; Hoffmann 2010; Buckley & Huey 2016). Our estimates of *D. melanogaster* adult male CT_max_ are consistent with previous reports (Gilchrist *et al.* 1997; Chown *et al.* 2009; Kellermann *et al.* 2012), and while we report novel findings on the adaptation of embryonic thermal tolerance, our results are not unprecedented. Coyne et al. (Coyne, Bundgaard & Prout 1983) reported a similar discrepancy in thermal adaptation between mobile and immobile life stages among populations of *Drosophila pseudoobscura*—pupal thermal tolerance, but not adult thermal tolerance, was higher in populations from warmer locations. The interplay of population genetic factors in natural populations of *D. melanogaster* suggest that this species harbors a high level of genetic diversity (Karasov, Messer & Petrov 2010) and that natural selection has led to allelic divergence among populations across the genome (Hoffmann & Weeks 2007; Fabian *et al.* 2012; Adrion *et al.* 2015). In light of these trends in population genomics, and the adaptive variation in embryonic thermal tolerance presented in this study, it seems probable that there is significant natural variation of upper thermal limits in *D. melanogaster* but that this variation may only be revealed in the embryonic and other immobile life stages.

It is important to note that laboratory selection experiments in *D. melanogaster*, *Escherichia coli*, and marine copepods (*Tigriopus californicus*) that imposed strong selection on thermal tolerance reported significant potential for adaptation of upper thermal limits, but the response to selection eventually plateaued after many generations, presumably when standing genetic diversity had been exhausted (Huey *et al.* 1991; Gilchrist *et al.* 1997; Gilchrist & Huey 1999; Rudolph *et al.* 2010; Kelly, Sanford & Grosberg 2012; Hangartner & Hoffmann 2015). Thus, there may likely be potential for adaptation of upper thermal limits, and in natural populations greater levels of standing genetic variation may be able to sustain adaptive responses to thermal selection.

This study characterizes thermal tolerance among populations that span a large portion of the *D. melanogaster* biogeographic range in the northern hemisphere, and while we present evidence of adaptation of embryonic thermal tolerance between temperate and tropical regions, the patterns of thermal adaptation are not consistent within each region. Tropical embryos sampled from locations with higher maximum habitat temperature (T_max_) showed higher thermal tolerances, yet temperate populations did not follow this trend. Why were there no observed differences in embryonic thermal tolerance among temperate populations when temperate sites spanned a broader range of thermal habitats than tropical populations? It is possible that gene flow between Vermont, Indiana, and North Carolina overwhelms local adaptation, but recent studies show evidence of adaptive divergence among *D. melanogaster* populations in eastern North America (Fabian *et al.* 2012; Bergland *et al.* 2016; Machado *et al.* 2016). Therefore, a more likely explanation is that seasonal fluctuations in the activity of temperate populations (Cogni *et al.* 2014), may limit the frequency at which temperate embryos encounter thermal selection. In addition, spatial and temporal microclimatic variability in temperate sites may provide more choices for females to lay their eggs at permissive temperatures (Allemand & David 1976; Dahlgaard, Hasson & Loeschcke 2001; Huey & Pascual 2009; Dillon *et al.* 2009).

We note that our data constitute thermal tolerances of multiple isofemale lines from each of the three temperate sites and one isofemale line from each of the five tropical sites. While we have not captured the full range of genetic variation within each tropical site, our data represent a broad sample of genetic diversity among tropical sites around the globe. Notably, the variance in thermal tolerance among all tropical genotypes was similar to the variance both within and among North American populations. However, there was no overlap in the confidence intervals of embryonic thermal tolerance between North American and tropical genotypes, whereas the confidence intervals of adult thermal tolerance were completely overlapping. Given that the tropical genotypes originated from geographically isolated locations (Table 1), we believe that these data reflect (1) selection for the maintenance of higher embryonic heat tolerance in the tropics and/or (2) convergent patterns of thermal adaptation across tropical populations. The positive correlation of embryonic thermal tolerance with maximum habitat temperature at tropical sites is a result that warrants further investigation. It remains to be determined the extent to which this pattern will hold when a greater sample of genetic diversity is surveyed within each topical population.

While thermal tolerance has been shown to be a complex quantitative trait in the adult and larval stages of *D. melanogaster* (Morgan & Mackay 2006; Sambucetti *et al.* 2013), the genetic basis of variation in embryonic thermal tolerance remains unresolved. We note that our reciprocal crossing design was not meant to be a full characterization of the genetic architecture of natural variation in embryonic thermal tolerance. Such an analysis would require a diallel crossing design among multiple isofemale lines in each population (Griffing 1956). Rather, our analysis was meant to test the potential role of maternal effects in our two most divergent genotypes (i.e. Chiapas [CH] vs. Vermont-12 [VT-12]). Because zygotic gene expression is inactive in early *D. melanogaster* embryos (0 – 1 h post-fertilization)(Tadros & Lipshitz 2009; Blythe & Wieschaus 2015), we predicted embryonic thermal tolerance to be determined by maternal factors, such as mRNAs and proteins, that are loaded into eggs. Contrary to this prediction, embryonic thermal tolerance in F1 progeny of crosses between Chiapas and Vermont-12 lines matched that of the Chiapas strain regardless of maternal genotype. This result suggests dominance of heat-tolerant alleles and not maternal effects as the basis of embryonic heat tolerance. Further, this suggests that either (1) the zygotic genome is being activated in embryos earlier than expected in response to heat shock (Graziosi *et al.* 1980), which would reveal adaptive variation in zygotic gene expression, or (2) that the effect is mediated at the level of the chromosomes, perhaps due to thermally-induced DNA damage (Yao & Somero 2012) that differentially affects different genotypes (Svetec *et al.* 2016). Either way, the unknown genetic basis of embryonic thermal tolerance warrants future study.

## Author’s Contributions

BL conceived the ideas and designed the methodology; BL, TG, and RS collected the data; BL analyzed the data; BL wrote the manuscript. All authors gave final approval for publication.

## Acknowledgements

We thank Kristi Montooth, Brandon Cooper, Cole Julick and the UC San Diego Drosophila Species Stock Center for providing fly stocks. We thank Sarah Howe for managing fly socks. We thank Emily Mikucki and Sarah Howe for their assistance in recording heat ramp rates. We thank Melissa Pespeni, Brandon Cooper, Kristi Montooth, Emily Mikucki, and two anonymous reviewers for helpful comments on this manuscript. This work was funded by the University of Vermont. The authors have no competing interests.

## Data Accessibility

Fly stock information is included in Table S1, including geographical coordinates of sampling locations, stock numbers, and thermal tolerance data.

---

**Supplemental Figure S1.**
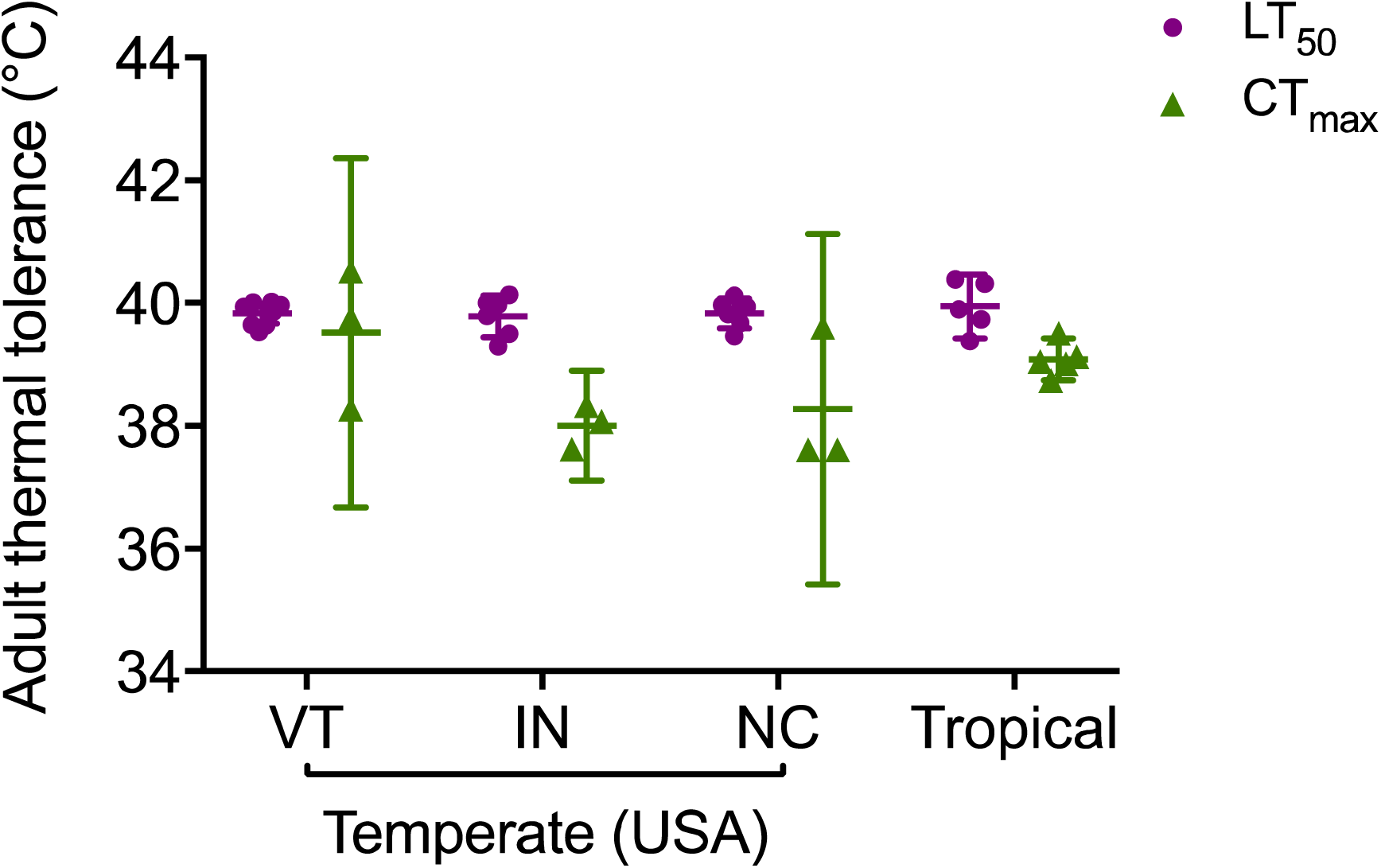
No significant difference in adult thermal tolerance (LT_50_ or CT_max_) among North American and tropical populations. There were no significant differences among collection sites in adult thermal tolerance as measured by LT_50_ (ANOVA, *F*_*3,20*_ = 0.3134, *P* = 0.8155) or CT_max_ (ANOVA, *F*_*3,9*_ = 2.378, *P* = 0.1375). Overall, adult CT_max_ values were lower than LT_50_ values across all sites (ANOVA, *F*_*1,31*_ = 44.73, *P* < 0.0001). Each point represents LT_50_ or CT_max_ for a single isofemale line. Error bars represent 95% confidence intervals and horizontal lines represent means among isofemale lines in each group.

**Supplemental Figure S2.**
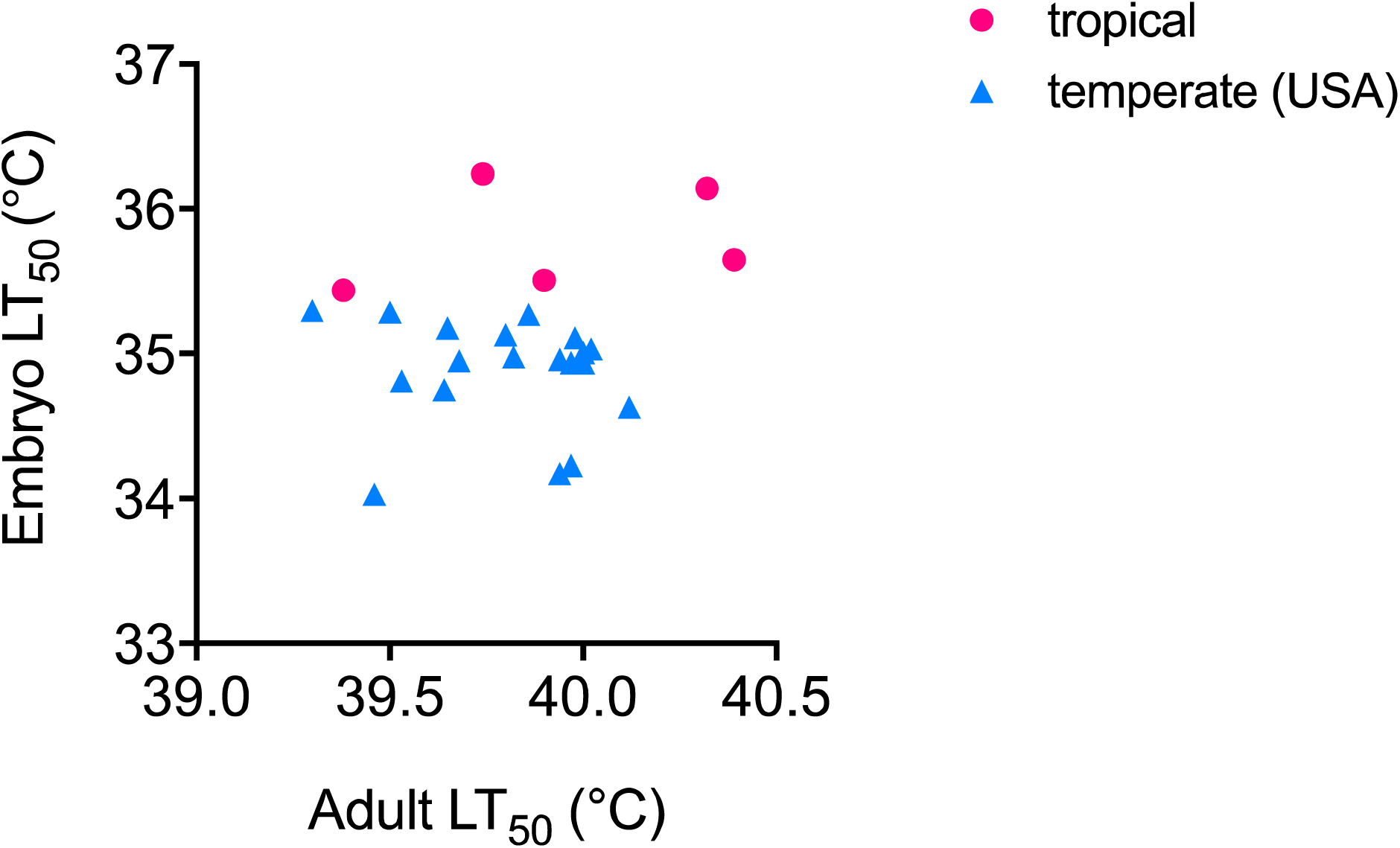
No significant relationship between adult and embryo thermal tolerance. Adult thermal tolerance and embryonic thermal tolerance were not correlated for either temperate (Least-squares linear regression, *R*^*2*^ = 0.015, *y* = -0.1973*x* + 42.73) or tropical lines (Least-squares linear regression, *R*^*2*^ = 0.09, *y* = 0.2664*x* + 25.15). Tropical isofemale lines are shown in pink circles and temperate isofemale lines are shown in blue triangles.

**Supplemental Figure S3.**
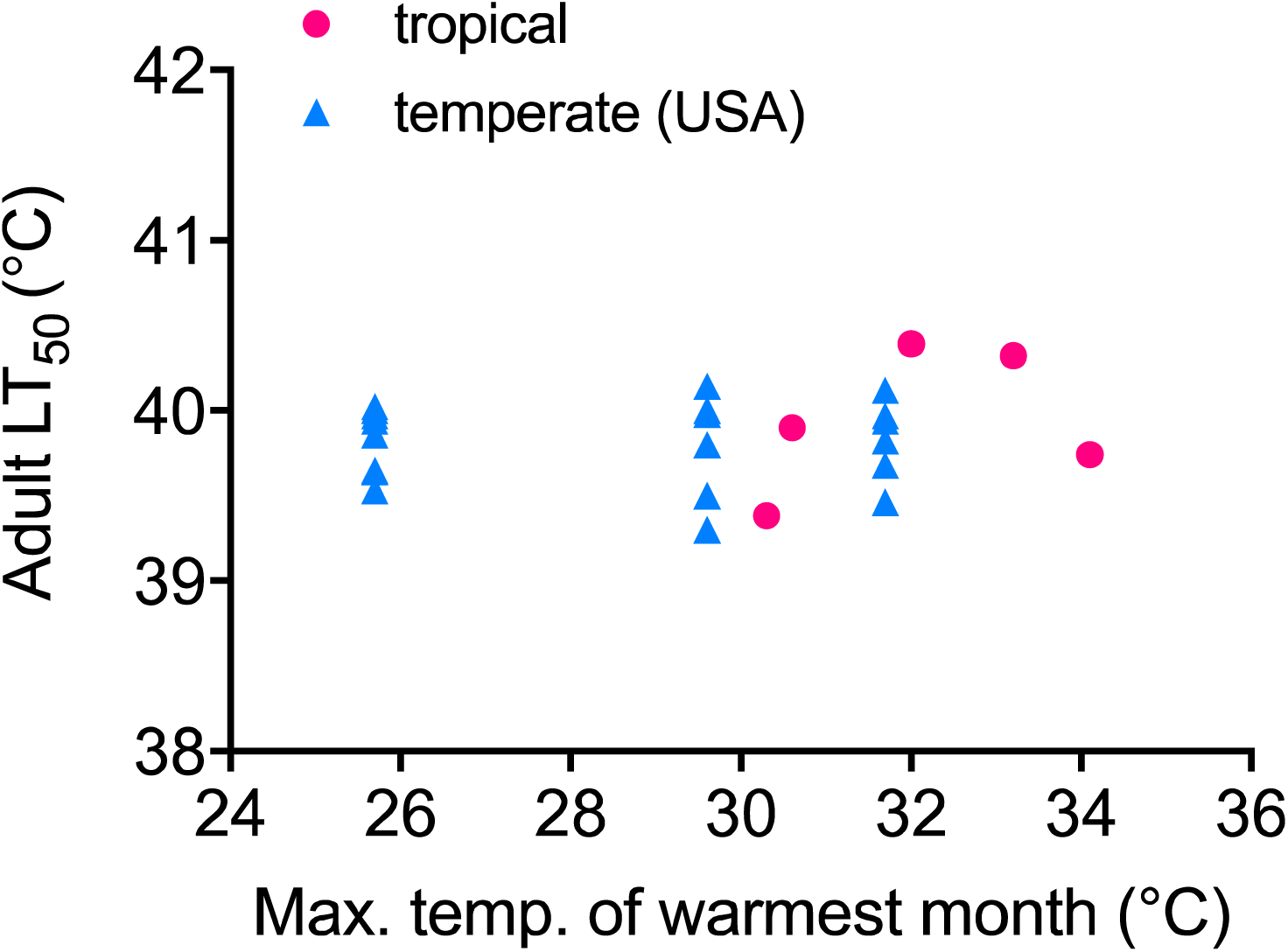
No significant relationship between adult thermal tolerance (LT_50_) and maximum habitat temperature (T_max_). Variation in adult thermal tolerance showed no significant correspondence to variation in T_max_ in either temperate (Least-squares linear regression, *R*^*2*^ = 0.004, *y* = -0.0005*x* + 39.83) or tropical regions (Least-squares linear regression, *R*^*2*^ = 0.14, *y* = 0.098*x* + 36.82). Tropical isofemale lines are shown in pink circles and temperate isofemale lines are shown in blue triangles.

**Supplementary Table S1:** Stock information, collection locale, year collected, and thermal tolerance data for isofemale lines used in this study.

| Stock No. | Locale | State/Country | Year | Lat. (°N) | Long. (°E) | Adult LT<sub>50</sub> (°C) | Embryo LT<sub>50</sub> (°C) |
| --- | --- | --- | --- | --- | --- | --- | --- |
| VTECK_2 | East Calais | Vermont, USA | 2011 | 44.37 | -72.43 | 39.65 | 35.18 |
| VTECK_4 | East Calais | Vermont, USA | 2011 | 44.37 | -72.43 | 40.02 | 35.03 |
| VTECK_5 | East Calais | Vermont, USA | 2011 | 44.37 | -72.43 | 39.64 | 34.75 |
| VTECK_8 | East Calais | Vermont, USA | 2011 | 44.37 | -72.43 | 39.53 | 34.81 |
| VTECK_9 | East Calais | Vermont, USA | 2011 | 44.37 | -72.43 | 39.86 | 35.27 |
| VTECK_10 | East Calais | Vermont, USA | 2011 | 44.37 | -72.43 | 39.94 | 34.17 |
| VTECK_12 | East Calais | Vermont, USA | 2011 | 44.37 | -72.43 | 39.97 | 34.23 |
| VTECK_14 | East Calais | Vermont, USA | 2011 | 44.37 | -72.43 | 40.00 | 35.01 |
| BEA_5 | Beasley Orchard | Indiana, USA | 2011 | 39.76 | -86.48 | 39.80 | 35.13 |
| BEA_16 | Beasley Orchard | Indiana, USA | 2011 | 39.76 | -86.48 | 39.98 | 35.11 |
| BEA_17 | Beasley Orchard | Indiana, USA | 2011 | 39.76 | -86.48 | 40.14 | - |
| BEA_21 | Beasley Orchard | Indiana, USA | 2011 | 39.76 | -86.48 | 40.00 | 34.94 |
| BEA_32 | Beasley Orchard | Indiana, USA | 2011 | 39.76 | -86.48 | 39.30 | 35.30 |
| BEA_36 | Beasley Orchard | Indiana, USA | 2011 | 39.76 | -86.48 | 39.50 | 35.29 |
| RFM_4 | Raleigh | North Carolina, USA | 2011 | 35.76 | -78.66 | 39.82 | 34.98 |
| RFM_6 | Raleigh | North Carolina, USA | 2011 | 35.76 | -78.66 | 39.68 | 34.95 |
| RFM_16 | Raleigh | North Carolina, USA | 2011 | 35.76 | -78.66 | 39.46 | 34.03 |
| RFM_19 | Raleigh | North Carolina, USA | 2011 | 35.76 | -78.66 | 40.12 | 34.63 |
| RFM_34 | Raleigh | North Carolina, USA | 2011 | 35.76 | -78.66 | 39.94 | 34.96 |
| RFM_48 | Raleigh | North Carolina, USA | 2011 | 35.76 | -78.66 | 39.97 | 34.94 |
| 14021-0231.22 | Chiapa de Corzo | Chiapas, Mexico | 2002 | 16.70 | -93.01 | 39.74 | 36.24 |
| 14021-0231.34 | Monkey Hill | St. Kitts | 2005 | 17.32 | -62.73 | 39.38 | 35.44 |
| 14021-0231.182 | Accra | Ghana | 2010 | 5.56 | -0.20 | 40.39 | 35.65 |
| 14021-0231.45 | Mumbai | Maharashtra, India | 2006 | 19.08 | 72.88 | 40.32 | 36.14 |
| 14021-0231.198 | Guam | Guam, USA | 2012 | 13.44 | 144.79 | 39.90 | 35.51 |

